# G-CSF associates with poor survival in cutaneous melanoma and promotes metastasis through coordinated effects on macrophages and tumor cells

**DOI:** 10.64898/2026.06.02.729497

**Authors:** Alicia Nieto-Valle, Lucía Barandalla-Revilla, Baltasar López-Navarro, Celia Barrio-Alonso, Alexandra de Francisco-López, José Antonio Avilés-Izquierdo, Verónica Parra-Blanco, Paloma Sánchez-Mateos, Rafael Samaniego

**Affiliations:** Unidad de Microscopía Confocal, Instituto de Investigación Sanitaria Gregorio Marañón, Madrid, Spain; Laboratorio de Inmuno-oncología, Instituto de Investigación Sanitaria Gregorio Marañón, Madrid, Spain; Departamento de Inmunología, Universidad Complutense de Madrid, Madrid, Spain; Unidad de Medicina y Cirugía Experimental, Instituto de Investigación Sanitaria Gregorio Marañón, Madrid, Spain; Servicio de Dermatología, Hospital General Universitario Gregorio Marañón, Madrid, Spain; Servicio de Anatomía Patológica, Hospital General Universitario Gregorio Marañón, Madrid, Spain

**Keywords:** G-CSF, CD114, melanoma, tumor microenvironment, tumor-associated macrophages, biomarker

## Abstract

Macrophage-melanoma interactions critically shape the tumor microenvironment, yet the cytokine networks driving this process remain incompletely understood. Among these, colony-stimulating factors—particularly granulocyte-macrophage colony-stimulating factor (GM-CSF) and granulocyte colony-stimulating factor (G-CSF or *CSF3*)—regulate myeloid cell behavior during cancer progression, although their specific roles in melanoma remain unclear. Using co-culture systems of melanoma and monocyte-derived macrophages, we found that GM-CSF-primed macrophages induced robust G-CSF secretion and concurrent *CSF3* upregulation in both cell compartments. Moreover, transcriptomic profiling of patient-derived tumor-associated macrophages (TAMs) confirmed elevated *CSF3* and *CSF3R* expression in metastatic melanoma. Multiplex immunofluorescence analysis of a stage II–IV primary melanoma cohort (n=84) revealed increased G-CSF and CD114 expression in TAMs and tumor cells from patients who subsequently developed metastasis. High G-CSF levels in either TAMs or melanoma cells emerged as an independent prognostic factor for shorter disease-free and overall survival (p < 0.001). Mechanistically, G-CSF activated STAT1/STAT3 signaling in melanoma cells and promoted proliferation and invasion in a CD114-dependent manner. Of note, G-CSF drove monocyte differentiation toward a distinct inflammatory macrophage state characterized by STAT1/STAT3 activation and a pro-invasive secretory profile. Furthermore, melanoma cells conditioned by G-CSF-differentiated macrophages displayed increased in vivo lung colonization and enriched transcriptional programs linked to invasion and proliferation. Collectively, these findings establish the G-CSF/CD114 axis as a potential clinically relevant driver of melanoma metastasis and identify G-CSF as a novel independent prognostic biomarker.

**HIGHLIGHTS:**

- Despite its extended clinical use, G-CSF role in human melanoma remains undefined.
- High G-CSF expression in primary melanomas correlates with poor patient survival.
- G-CSF promotes melanoma cell invasion and proliferation through CD114 signaling.
- G-CSF drives macrophage differentiation toward a pro-tumoral phenotype.
- Melanoma cells conditioned by G-CSF-primed macrophages are more metastatic.

## INTRODUCTION

Tumor-associated macrophages (TAMs) are key innate immune cells in the tumor microenvironment (TME) due to their multiple pro-tumorigenic functions, and their high infiltration is consistently associated with poor clinical outcomes in many cancers [1,2]. Macrophage identity and functional heterogeneity is tightly regulated by the surrounding environment, as TAMs exhibit extraordinary plasticity and can acquire a broad spectrum of activation and functional states. Single-cell and lineage-tracing technologies have exposed how TAM biology is inherently complex, with the co-existence of distinct subpopulations with a differential ontogeny, spatial tissue localization, and cancer stage-dependent functions that evolve over time [2]. Recent omic reports unveil the multidimensional expression patterns that link with functional TAM diversity and can hold prognostic value [1,3].

Recognized as a hallmark of cancer, chronic inflammation is repeatedly shown to promote cancer initiation, growth, progression and metastasis [4], and it is increasingly evident that pro-inflammatory macrophages may sustain cytotoxic pressure that selects for more malignant and immune-resistant tumor clones [5,6]. In primary cutaneous melanoma, while we found no significant association between TAM density, size, or location with patient survival, adverse prognosis did correlate with a secretory molecular signature that aligned with an “inflammatory cytokine-enriched” TAM phenotype [3,7]. Furthermore, we recently reported that the expression of granulocyte macrophage colony-stimulating factor (GM-CSF), but not macrophage colony-stimulating factor (M-CSF), advocates for a pro-inflammatory milieu in macrophage-tumor environment that promotes cancer invasion and metastasis and correlates with poor patient outcome [8].

Along with GM-CSF and M-CSF, granulocyte colony-stimulating factor (G-CSF) is a hematopoietic growth factor that regulates myeloid cell behavior during inflammatory responses and emergency myelopoiesis [9,10]. G-CSF receptor (G-CSFR or CD114) is expressed in a wide variety of immune and non-immune cells, including myeloid lineage, namely neutrophils and monocytes, myeloid progenitors, endothelial cells and tumor cells, with controversial reports of its expression on lymphocytic cells [11]. Although extensively used in clinical practice to treat chemotherapy-induced neutropenia [12], solid tumors may aberrantly produce G-CSF and express its receptor and to subsequently modulate immune responses of incoming macrophages, myeloid-derived suppressor cells (MDSCs) or regulatory T cells, and promote tumor stemness, proliferation or migration [13]. Furthermore, tumor-derived G-CSF may mediate the systemic deviation of hematopoiesis and acceleration of myelopoiesis, resulting in leukocytosis and elevated G-CSF blood levels, as it has been reported in some melanoma patients [14]. Remarkably, healthy donors treated with G-CSF for standard hematopoietic stem cell mobilization present the typical mononuclear and polymorphonuclear MDSCs phenotypes described in cancer patients [15]. This uncovers a complex and multidimensional pro-tumoral mechanism of G-CSF by which it directly enhances tumor cell viability, proliferation and invasion, and modulates immune populations both locally and systemically.

G-CSF-secreting tumors are highly aggressive and directly linked with secondary metastasis, worse prognosis and low survival rates, while the dynamic changes caused by G-CSF in the TME can be used as markers of early disease progression and therapeutic response [13]. The expression of both G-CSF and G-CSFR, encoded by *CSF3* and *CSF3R* genes, respectively, has been evaluated across several tumors to elucidate their potential association with prognosis. In melanoma, while public datasets like TCGA suggest a positive correlation between *CSF3R* expression and patient survival [16], clinical data conversely associate elevated G-CSF plasma levels with resistance to anti-CTLA-4 immunotherapy [17]. Despite this conflicting evidence, no reports have specifically addressed the expression of G-CSF/G-CSFR in the human melanoma microenvironment. In this study, we evaluated the expression of G-CSF and its receptor in a cohort of primary melanoma samples and assessed their potential association with metastatic progression. Furthermore, we investigated the functional effects of G-CSF on both melanoma cells and monocytes, as well as its possible contribution to the mechanisms underlying melanoma metastasis.

## MATERIALS AND METHODS

### Study cohort and selection criteria

Patient samples were collected following the approval of the Hospital General Universitario Gregorio Marañón ethics committee, and informed consent was obtained for each patient. A formalin-fixed and paraffin-embedded (FFPE) primary cutaneous melanoma cohort of 84 patients was used, with a > 2 mm Breslow thickness, American Joint Committee on Cancer pathological staging II–IV, and a median follow-up of 67.5 months, as previously described [18]. This cohort included 34 samples from patients who were disease-free for at least 10 years of follow-up (non-metastasizing primary melanomas) and 50 clinically aggressive samples developing distant metastasis (metastasizing primary melanomas, with 30/50 related deaths). Six patients at stage IV were excluded from disease-free survival (DFS) analysis, but not from overall survival (OS).

### Multicolor fluorescence confocal microscopy

FFPE samples were deparaffinized, rehydrated, and unmasked by steaming in 10 mM sodium citrate buffer pH 9.0 (Dako) for 7 minutes. Slides were blocked with 5 μg/mL human immunoglobulins solved in blocking serum-free medium (Dako) for 30 min and then sequentially incubated with 5–10 μg/mL primary antibodies (supplementary Table S1) and appropriate fluorescent secondary antibodies (Jackson Immunoresearch), as previously described [18]. Samples were imaged with a SPE confocal microscope using the glycerol-immersion ACS APO x20/NA 0.60 objective (Leica Microsystems). Single-cell quantification was performed at 3-5 20x fields, and Mean Fluorescence Intensity (MFI) of proteins obtained at manually depicted tumor cells or at segmented CD68^+^ TAMs using the ‘analyze particle’ plugin of ImageJ2 software, as previously described [19]. For in vivo lung colonization assays, NG2^+^ melanoma micrometastases were microscopically measured at multiple histologic frozen sections.

### Monocyte isolation and cell culture

Biopsied primary melanomas (n= 4) were homogenized and digested into single-cell suspensions (Tumor Dissociation Kit, Miltenyi Biotec) and peripheral blood mononuclear cells were isolated from buffy coats from melanoma patients or healthy donors over a Lymphoprep™ gradient (STEMCELL Technologies). Both TAMs and monocytes were purified by magnetic cell sorting using anti-CD14 tagged microbeads (Miltenyi Biotech). For in vitro priming, healthy donor monocytes were cultured at 0.5 x 10^6^ cells/mL for 7 days containing recombinant human M-CSF (rhM-CSF, 10 ng/mL, Immunotools), recombinant human GM-CSF (rhGM-CSF, 10 ng/mL, Immunotools) or recombinant human G-CSF (rhG-CSF, 25 ng/mL, Immunotools) to generate M-Macs, GM-Macs or G-Macs, respectively. GM/G-Macs and M/G-Macs macrophages were obtained by combining GM-CSF or M-CSF with G-CSF during the 7 days of priming. Cytokines were added every two days. Due to the lower baseline viability, at day 3 G-Macs and Mo-Macs cells were detached, counted, and re-seeded at equal cell densities. For in vitro staining, macrophages were seeded at day 6 at a density of 10^5^ cells/well in poly-L-lysine (0.01%) coated coverslips for 2 hours and fixed with 1% formaldehyde for 10 minutes at room temperature. Afterwards, cells were permeabilized and stained with phalloidin-iFluor555 (Abcam) and anti-CD14-FITC antibody (supplementary Table S1). All cells, including the melanoma cell lines BLM, A375, and SK-MEL-103 [18] were cultured in RMPI-1640 medium (Gibco) supplemented with 10% fetal calf serum (FCS, Sigma-Aldrich) and a mix of antibiotics (8 µg/mL gentamicin, 10 µg/mL cloxacillin, and 10 µg/mL ampicillin).

### In vitro measurements

Melanoma cells were co-cultured with monocytes or primed macrophages at different melanoma:myeloid cell ratios. For secretion analysis, conditioned media (CM) were collected after 72 hours of a 1:4 ratio culture for Quantibody multiplex array (RayBiotech) or 48 hours for specific G-CSF ELISA kit (R&D Systems). For transcriptomic analysis, TCs and monocytes/macrophages were cultured at 1:2 ratio and processed after 24 hours, separating both cell types with anti-CD14 microbeads. Total RNA was processed and sequenced at BGI (Shenzhen) using the DNBseq-G400 platform. For protein expression analysis of the G-CSF receptor, whole-cell lysates were subjected to western blotting with the indicated antibodies (supplementary Table S1). For signaling kinetics, human monocytes (2.5 × 10^6^) or melanoma cell lines (1 × 10^6^) were starved in serum-free medium for 1 hour before supplementation with distinct G-CSF concentrations for 15 or 30 minutes. Cell lysates were directly prepared using 1x Laemmli sample buffer. For surface marker staining and flow cytometry, cell suspensions were blocked with human immunoglobulins and stained with conjugated antibodies for 25 minutes at 4°C (supplementary Table S1). Data were acquired on a CytoFLEX flow cytometer and analyzed with Kaluza™ software (Beckman Coulter).

### Transcriptomic analysis

Differentially-expressed genes (DEGs) were assessed by using DESeq2 algorithm with parameters

|log_2_ (fold change)| > 2 and adjusted p value < 0.05. For interactome analysis, pairs of ligands and receptors listed in the lr_network of NicheNet tool [20] were evaluated, and interactions were considered active when both ligand and receptor were upregulated DEGs in the two interacting cells. All active ligand-receptor interactions were summarized in circos plots using the circlize R package [21]. To analyze each ligand regulation potential, we performed enrichment analysis on lists of the top 50 most regulated genes according to the ligand_target_matrix of NicheNet (supplementary Table S4), [20]. Gene Set Enrichment Analysis (GSEA) was performed using the clusterProfiler R package [22] to evaluate the coordinated regulation of defined gene collections, including the hallmark gene sets from the Molecular Signature Database (MSigDB) [23]. Prior to analysis, genes were ranked according to the Wald statistic (stat value) obtained from DESeq2. Transcription factor activities were inferred using the DoRothEA (Discriminant Regulon Expression Analysis) resource based on expression changes of their downstream target genes [24]. RNA sequencing (RNA-seq) data was deposited at Gene Expression Omnibus under accession numbers GSE171277, GSE242674 and GSE(pending).

### Viability, invasion and proliferation assays

Fresh monocytes were seeded at 2.5 x 10^5^ cells/well with none, M-CSF, GM-CSF and/or G-CSF in 48-well plates for 9 days and cell viability was assessed every 3 days using the CCK-8 kit (MedChemExpress) following the manufacturer’s instructions. For invasion assays, BLM and SK-MEL-103 spheroids were produced by seeding 10^4^ cells per well in a 96-well U-bottom plate (Thermo Fisher Scientific) for 5–7 days, embedded in 2 mg/mL collagen type-I (STEMCELL Technologies) gels and allowed to invade for 72 hours in the presence of serum-deprived media ± G-CSF, and ± 25 μg/mL of neutralizing antibodies for human CD114 or isotype-control antibodies (Biolegend; Supplementary Table S1). Tumor cells and/or mMonocytes/macrophages were first co-cultured at 1:4 ratio for 48 hours in 10% FCS medium, and then media were replaced by serum-free medium for 24 additional hours to obtain the CM used in invasion assays. Quantifications were performed as previously described [18]. For proliferation assays, melanoma cells were seeded at a density of 10^5^ cells/well in poly-L-lysine (0.01%) coated coverslips with 1% FCS medium supplemented with rhG-CSF at different concentrations. After two days of culture for A375 and SK-MEL-103, and four days in the case of BLM, cells were incubated with 10 μg/ml 5-bromo-2’-deoxyuridine (BrdU, Sigma-Aldrich) for 1.5 hours and then fixed with 1% formaldehyde for 10 minutes at room temperature. For BrdU incorporation staining, cells were permeabilized and stained with 5μg/ml anti-BrdU (R&D Systems) and a conjugated secondary antibody. DAPI-stained nuclei were segmented and BrdU MFI was assessed using ImageJ2, as previously described [18].

### In vivo human melanoma models

NOD/SCID/IL-2Rγ −/− mice (NSG, The Jackson Laboratory) were maintained under specific pathogen-free conditions. For lung colonization experiments, male or female mice were used at the age of 6-8 weeks (∼23 g). After 72 hours of co-culture with macrophages (1:4 ratio, melanoma:macrophage), conditioned melanoma cells —named as TC (Mac)— were administered via intravenous tail vein injection at a concentration of 0.5 × 10^6^ tumor cells suspended in 150 μL of PBS. At day 6 (BLM) or 13 (A375), lungs were extracted for quantitative histologicalanalyss. All procedures were approved by the IiSGM animal care/use and Comunidad de Madrid committees (Proex: 296.4/22 and 132.3/25).

### Statistical analyses

Kaplan–Meier curves were used to analyze the correlation between proteins expression and patient DFS and OS, using Youden’s index to determine where the cut-off point was equally specific and sensitive. The Cox regression method (univariate and multivariate) was used to identify independent prognostic variables and Mann–Whitney tests to evaluate the association with clinicopathological features. ANOVA analyses, paired and unpaired t-test and log-rank analyses were also used in this study (GraphPad software and R), as indicated; p< 0.05 was considered statistically significant.

## RESULTS

### Interaction between GM-CSF-primed macrophages and melanoma cells activates G-CSF/CD114 axis

We recently demonstrated that the microenvironment of metastasizing primary melanomas is preferentially enriched in GM-CSF, whereas M-CSF expression remains constitutive and independent of tumor progression [8]. To further investigate the crosstalk between macrophages and melanoma cells, we established an in vitro co-culture system of BLM melanoma cells with monocyte-derived macrophages previously primed with either anti-inflammatory M-CSF or pro-inflammatory GM-CSF. We evaluated the secretory profile through a cytokine array panel of 162 growth factors and interleukins (Supplementary Table S2), as previously done for chemokines [18]. A subset of cytokines was markedly upregulated following macrophage–melanoma interaction (Figure 1A), with G-CSF emerging as prominently enriched in both experimental settings (Figure 1B). These findings were extended and validated by ELISA, confirming that G-CSF secretion was significantly higher in BLM and A375 melanoma co-cultures with GM-CSF-primed macrophages (GM-Macs) compared to M-CSF-primed macrophages (M-Macs) or untreated monocytes (Figure 1C).

**Figure 1.**
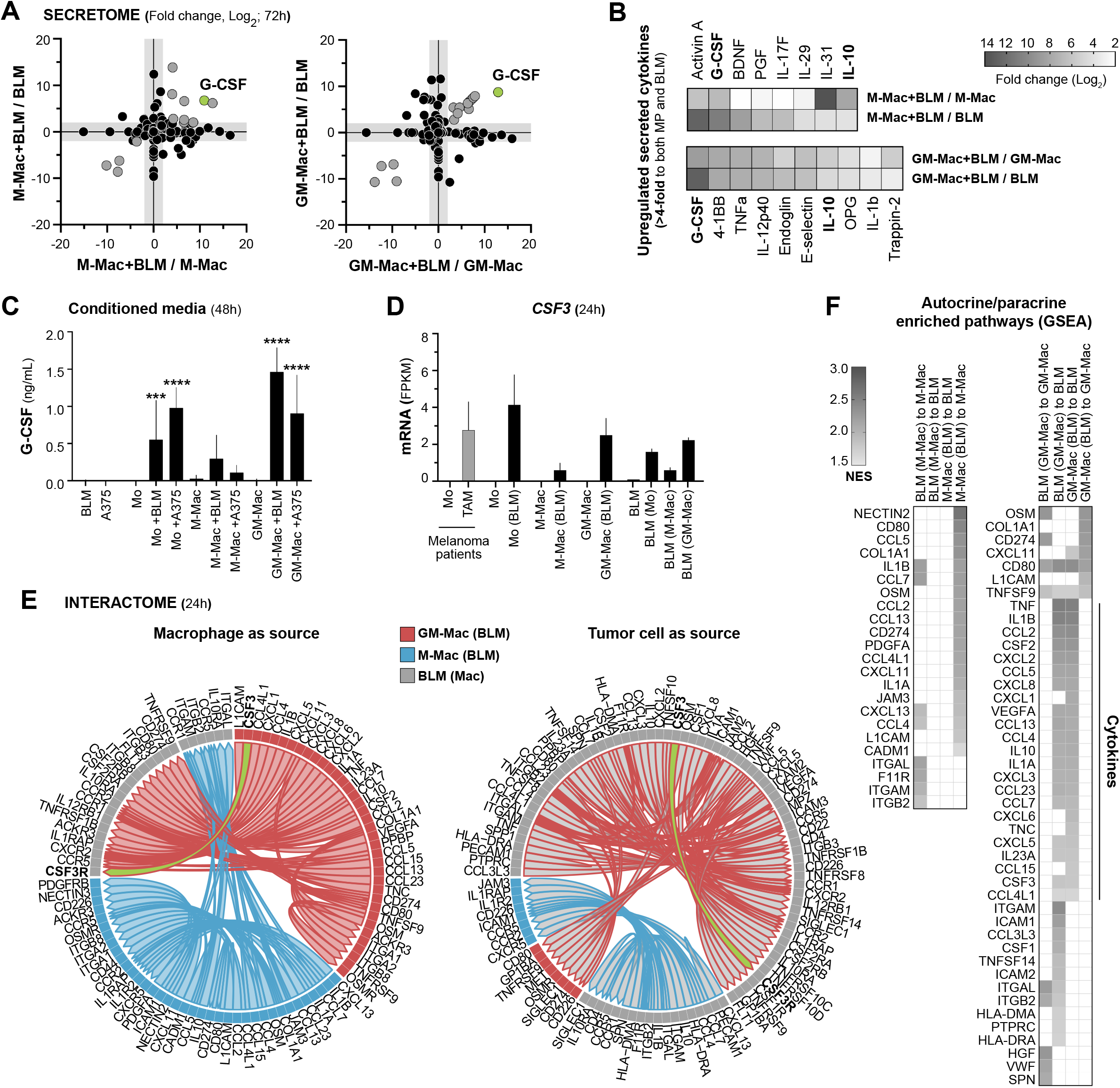
Co-culture of monocytes/macrophages and melanoma cells induces G-CSF/CD114 axis. Monocytes, M-CSF-primed macrophages (M-Macs) and GM-CSF-primed macrophages (GM-Macs) were co-cultured with BLM or A375 melanoma cells for distinct transcriptomic (24 hours) and secretomic (48 or 72 hours) analyses. **(A)** Secretomic analysis (163 cytokines) of 72 hours co-cultured supernatants comparing M-Mac+BLM or GM-Mac+BLM with unexposed macrophages (x-axis) or BLM (y-axis) cells. Fold changes (Log_2_) are shown (n = 2 donors). **(B)** Heatmap of most upregulated secreted cytokines after co-culture of BLM with M-Macs or GM-Macs (fold change > 4 to both unexposed macrophages and melanoma cells, are shown). **(C)** G-CSF secretion assessed at monocytes/macrophages ± melanoma cells conditioned media at 48 hours. Mean ± standard deviation (SD) values (n = 4-5 donors), are shown. Significant differences to the respective untreated controls are indicated (***p < 0.001; ****p < 0.0001, paired t-test). **(D)** *CSF3* mRNA levels on both monocytes/macrophages and tumor cells isolated 24 hours after co-culture, as well as monocytes and primary melanoma TAMs isolated from stage-IV melanoma patients. Mean ± SEM values (n = 3 donors), are shown. **(E)** Interactome analysis based on the concurrent upregulation of ligand-receptor genes in macrophages and BLM cells after 24 hours co-culture. **(F)** Gene set enrichment analyses (GSEA) for autocrine and paracrine potential target genes regulated by distinct upregulated ligands in macrophage-melanoma co-culture. Normalized Enrichment Score (NES) is shown for significantly enriched pathways (adjusted p < 0.05).

To identify the cellular source(s) of G-CSF, RNA sequencing (RNA-seq) was performed on macrophages and melanoma TCs isolated after co-culture. *CSF3* mRNA expression was upregulated in both cell types, although induction was less pronounced in the M-CSF-priming condition (Figure 1D). Expression analysis of circulating monocytes and TAMs isolated from stage-IV melanoma patients revealed minimal *CSF3* transcription in blood monocytes but elevated levels in TAMs, suggesting that monocytes acquire a G-CSF-secreting phenotype within the TME (Figure 1D). To further characterize the signaling circuitry established between macrophages and melanoma cells, we conducted an interactome analysis based on the concurrent upregulation of ligand-receptor pairs. Herein, we identified several newly established chemokine/cytokine axes in co-cultured cells, including a *CSF3*-driven axis originating from both melanoma cells and macrophages and targeting *CSF3R* expressed by melanoma cells in the GM-CSF-primed context (Figure 1E). Subsequently, to evaluate the autocrine and paracrine regulatory potential of these distinct ligands in our macrophage-melanoma co-culture samples, we performed GSEA on defined gene sets comprising the top 50 target genes with the highest NicheNet probability scores (Supplementary Table S4) [20] to identify actively regulated pathways (Figure 1F). Collectively, these findings demonstrate that the interaction between GM-Macs and melanoma cells induces an extensive autocrine/paracrine network of active ligands, including various cytokines and *CSF3*. Along with our previous results on the correlation with a cytokine-enriched inflammatory TAM phenotype [8,18], this result may reflect a synergistic and/or mediating role for GM-CSF in potentiating the G-CSF/CD114 axis in human melanoma.

### G-CSF and G-CSFR expression in primary melanoma tissues and association with patient survival

To investigate the clinical relevance of G-CSF in cutaneous melanoma, we first assessed its expression at single-cell resolution using multiplex immunofluorescence. An independent cohort of 84 stage II–IV paraffin-embedded primary cutaneous melanomas with annotated clinical follow-up was analyzed. Patients were stratified according to the development of distant metastasis within 10 years following tumor resection. G-CSF and its receptor CD114 were significantly enriched in both TAMs and TCs from metastasizing primary melanomas compared to non-metastasizing tumors (Figure 2A, B). Furthermore, CD114 expression in TCs was enriched in deeper lesions according to Breslow thickness (Mann-Whitney, p < 0.001), suggesting its potential association with primary tumor growth (Supplementary Table S3). To evaluate their prognostic value, G-CSF and CD114 expression levels within both cell types were used to independently stratify the patient cohort for Kaplan–Meier survival analyses. G-CSF expression in both cell compartments, but not CD114 expression, was significantly associated with reduced disease-free survival (log-rank p < 0.0001) and overall survival (p < 0.0001) (Figure 2C, D; Supplementary Figure S1C, D).

**Figure 2.**
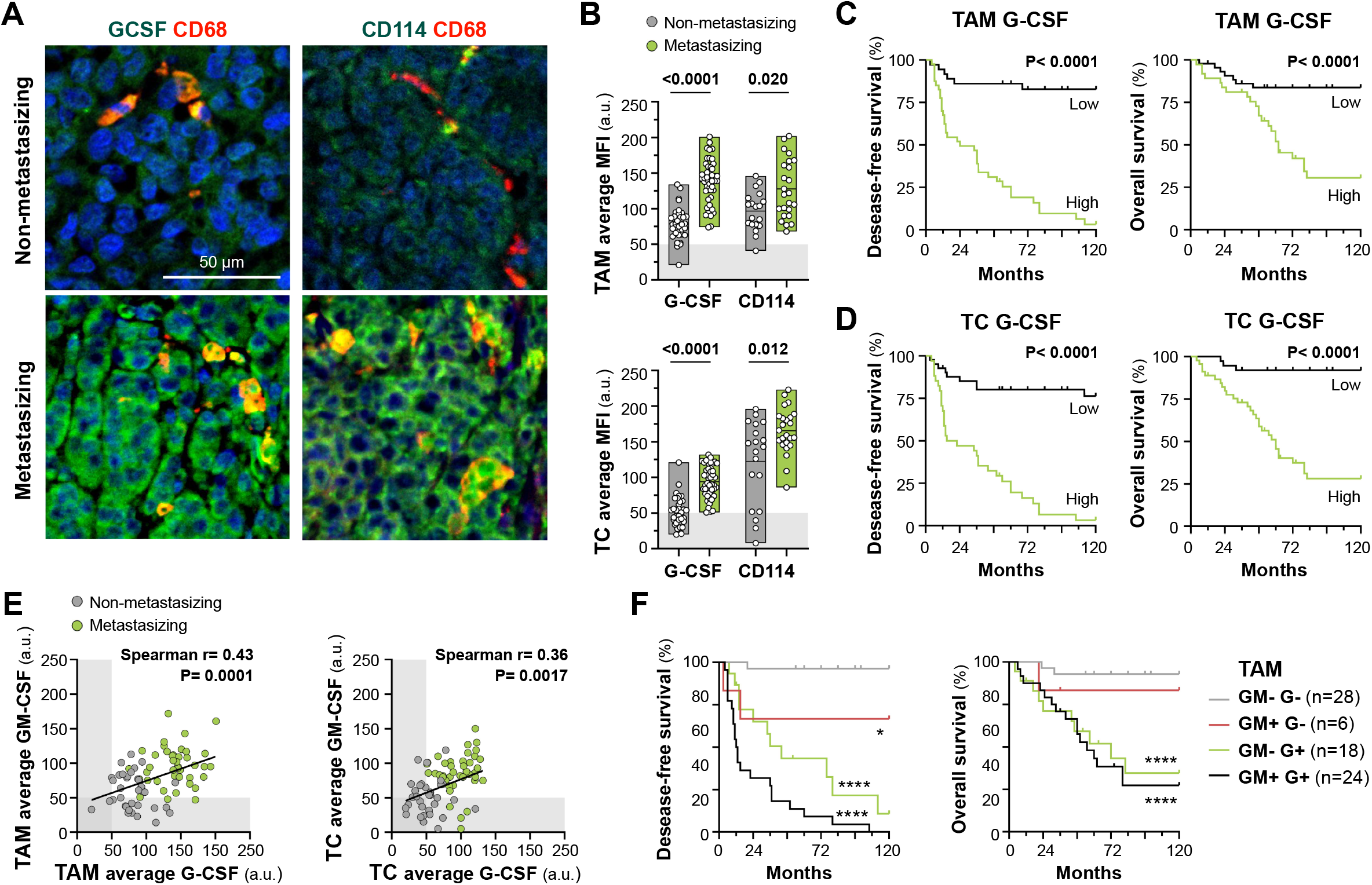
Expression of G-CSF and its receptor CD114 in human primary melanomas and correlation with patient survival. **(A)** Representative FFPE sections of non-metastasizing and metastasizing primary melanoma samples co-stained for the pan-macrophage marker CD68 (red), the proteins of interest G-CSF and CD114 (green) and Dapi (blue), as indicated. **(B)** Single-cell quantification of G-CSF and CD114 in CD68^+^ TAMs and TCs. Plots show the average MFIs (arbitrary units, a.u.) of analyzed proteins for each tumor sample. Mann–Whitney p values are shown. **(C, D)** Disease-free and overall survival 10-year Kaplan–Meier curves for G-CSF expression in TAMs (C; cut-off value: 100 a.u.) and TCs (D; cut-off value: 80 a.u.). Log-rank p value*s* are shown. **(E)** Correlative expression of GM-CSF and G-CSF in TAMs and TCs for each tumor sample. Spearman’s correlation analyses are shown. **(F)** Disease-free and overall survival 10-year Kaplan–Meier curves for GM-CSF and G-CSF expression in TAMs, categorized by high (+) or low (-) expression of both cytokines, as indicated. Log-rank p value*s* relative to double negative samples are shown (*p < 0.05; ****p < 0.0001). Scale bar, 50 μm.

As suggested in the previous section, GM-CSF might be playing a synergistic and/or mediating role in potentiating the G-CSF/CD114 axis. Indeed, G-CSF and GM-CSF expression levels in TAMs and TCs positively correlated in primary melanomas (Spearman r = 0.4, p < 0.001) (Figure 2E). Kaplan-Meier survival analysis revealed that patients with low or undetectable expression of both cytokines in both TAMs and TCs exhibited a favorable prognosis, while double-positive samples reached the worst prognostic rate for both DFS and OS (Figure 2F; Supplementary Figure S1E). Of note, the presence of G-CSF or GM-CSF alone in TCs was sufficient to predict poor clinical outcomes, while in TAMs only G-CSF alone correlated with shorter DFS and OS (p < 0.0001). Moreover, multivariate Cox regression analysis further identified TAM G-CSF expression as an independent prognostic factor for both DFS (Hazard Ratio = 19.9; p < 0.001) and OS (HR = 9.1; p = 0.002), after adjustment for age, sex, Breslow thickness, tumor stage, and TAM GM-CSF expression (Table 1). In the case of melanoma TCs, both G-CSF and GM-CSF expression levels were independent prognostic factors for both DFS and OS (Table 1). Elevated G-CSF levels were preferentially found in tumors enriched in pro-inflammatory cytokine-producing TAMs (Supplementary Figure S1A), independently of their overall abundance (Supplementary Figure S1D), and did not correlate with accumulations of neutrophils or MDSCs (Supplementary Figure S1D), which are scarce in primary human melanomas [7]. Together, our multiplex analyses demonstrate that metastasizing primary melanomas are characterized by a G-CSF-enriched microenvironment, supporting a potential pro-metastatic role for this cytokine through its action on both macrophages and melanoma cells, which express the cognate receptor.

**Table 1.**
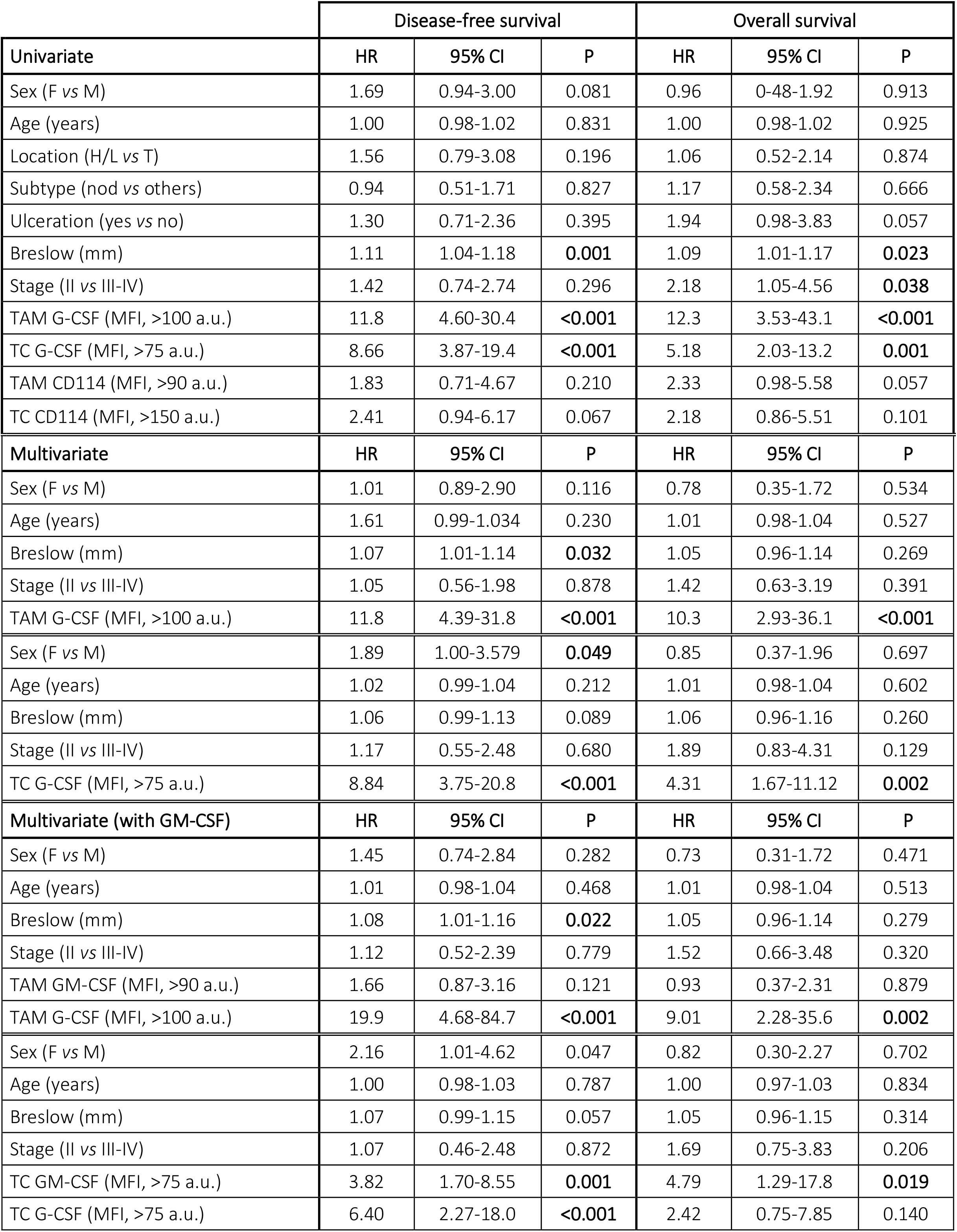
Univariate and multivariate Cox regression analysis.

|  | Disease-free survival |  |  | Overall survival |  |  |
| --- | --- | --- | --- | --- | --- | --- |
| Univariate | HR | 95% CI | P | HR | 95% CI | P |
| Sex (F vs M) | 1.69 | 0.94-3.00 | 0.081 | 0.96 | 0-48-1.92 | 0.913 |
| Age (years) | 1.00 | 0.98-1.02 | 0.831 | 1.00 | 0.98-1.02 | 0.925 |
| Location (H/L vs T) | 1.56 | 0.79-3.08 | 0.196 | 1.06 | 0.52-2.14 | 0.874 |
| Subtype (nod vs others) | 0.94 | 0.51-1.71 | 0.827 | 1.17 | 0.58-2.34 | 0.666 |
| Ulceration (yes vs no) | 1.30 | 0.71-2.36 | 0.395 | 1.94 | 0.98-3.83 | 0.057 |
| Breslow (mm) | 1.11 | 1.04-1.18 | **0.001** | 1.09 | 1.01-1.17 | **0.023** |
| Stage (II vs III-IV) | 1.42 | 0.74-2.74 | 0.296 | 2.18 | 1.05-4.56 | **0.038** |
| TAM G-CSF (MFI, >100 a.u.) | 11.8 | 4.60-30.4 | **<0.001** | 12.3 | 3.53-43.1 | **<0.001** |
| TC G-CSF (MFI, >75 a.u.) | 8.66 | 3.87-19.4 | **<0.001** | 5.18 | 2.03-13.2 | **0.001** |
| TAM CD114 (MFI, >90 a.u.) | 1.83 | 0.71-4.67 | 0.210 | 2.33 | 0.98-5.58 | 0.057 |
| TC CD114 (MFI, >150 a.u.) | 2.41 | 0.94-6.17 | 0.067 | 2.18 | 0.86-5.51 | 0.101 |
| Multivariate | HR | 95% CI | P | HR | 95% CI | P |
| Sex (F vs M) | 1.01 | 0.89-2.90 | 0.116 | 0.78 | 0.35-1.72 | 0.534 |
| Age (years) | 1.61 | 0.99-1.034 | 0.230 | 1.01 | 0.98-1.04 | 0.527 |
| Breslow (mm) | 1.07 | 1.01-1.14 | **0.032** | 1.05 | 0.96-1.14 | 0.269 |
| Stage (II vs III-IV) | 1.05 | 0.56-1.98 | 0.878 | 1.42 | 0.63-3.19 | 0.391 |
| TAM G-CSF (MFI, >100 a.u.) | 11.8 | 4.39-31.8 | **<0.001** | 10.3 | 2.93-36.1 | **<0.001** |
| Sex (F vs M) | 1.89 | 1.00-3.579 | **0.049** | 0.85 | 0.37-1.96 | 0.697 |
| Age (years) | 1.02 | 0.99-1.04 | 0.212 | 1.01 | 0.98-1.04 | 0.602 |
| Breslow (mm) | 1.06 | 0.99-1.13 | 0.089 | 1.06 | 0.96-1.16 | 0.260 |
| Stage (II vs III-IV) | 1.17 | 0.55-2.48 | 0.680 | 1.89 | 0.83-4.31 | 0.129 |
| TC G-CSF (MFI, >75 a.u.) | 8.84 | 3.75-20.8 | **<0.001** | 4.31 | 1.67-11.12 | **0.002** |
| Multivariate (with GM-CSF) | HR | 95% CI | P | HR | 95% CI | P |
| Sex (F vs M) | 1.45 | 0.74-2.84 | 0.282 | 0.73 | 0.31-1.72 | 0.471 |
| Age (years) | 1.01 | 0.98-1.04 | 0.468 | 1.01 | 0.98-1.04 | 0.513 |
| Breslow (mm) | 1.08 | 1.01-1.16 | **0.022** | 1.05 | 0.96-1.14 | 0.279 |
| Stage (II vs III-IV) | 1.12 | 0.52-2.39 | 0.779 | 1.52 | 0.66-3.48 | 0.320 |
| TAM GM-CSF (MFI, >90 a.u.) | 1.66 | 0.87-3.16 | 0.121 | 0.93 | 0.37-2.31 | 0.879 |
| TAM G-CSF (MFI, >100 a.u.) | 19.9 | 4.68-84.7 | **<0.001** | 9.01 | 2.28-35.6 | **0.002** |
| Sex (F vs M) | 2.16 | 1.01-4.62 | 0.047 | 0.82 | 0.30-2.27 | 0.702 |
| Age (years) | 1.00 | 0.98-1.03 | 0.787 | 1.00 | 0.97-1.03 | 0.834 |
| Breslow (mm) | 1.07 | 0.99-1.15 | 0.057 | 1.05 | 0.96-1.15 | 0.314 |
| Stage (II vs III-IV) | 1.07 | 0.46-2.48 | 0.872 | 1.69 | 0.75-3.83 | 0.206 |
| TC GM-CSF (MFI, >75 a.u.) | 3.82 | 1.70-8.55 | **0.001** | 4.79 | 1.29-17.8 | **0.019** |
| TC G-CSF (MFI, >75 a.u.) | 6.40 | 2.27-18.0 | **<0.001** | 2.42 | 0.75-7.85 | 0.140 |

### G-CSF promotes melanoma cell proliferation and invasion

Given the widespread expression of CD114 in melanoma cells across primary tumors, we next investigated whether G-CSF might directly modulate melanoma cell functions. CD114 expression was confirmed by immunoblotting in A375, SK-MEL-103, and BLM melanoma cell lines (Figure 3A). We then examined downstream signaling pathways known to be activated by G-CSF. Exposure to rhG-CSF induced phosphorylation of STAT1 and STAT3, but not STAT5, under the tested conditions, indicating that the receptor is functional and capable of triggering intracellular signaling cascades relevant to melanoma cell biology (Figure 3A). To assess functional consequences, we performed proliferation and 3D-collagen invasion assays. Melanoma cells cultured with increasing concentrations of rhG-CSF exhibited enhanced proliferation, as measured by BrdU incorporation, across all three cell lines (Figure 3B). In parallel, BLM and SK-MEL-103 spheroids embedded in collagen matrices exhibited dose-dependent increases in invasive capacity upon G-CSF treatment. This effect was abrogated by a neutralizing anti-CD114 antibody, confirming receptor specificity (Figure 3C). These data demonstrate that G-CSF directly enhances melanoma cell proliferation and invasion through CD114-dependent signaling.

**Figure 3.**
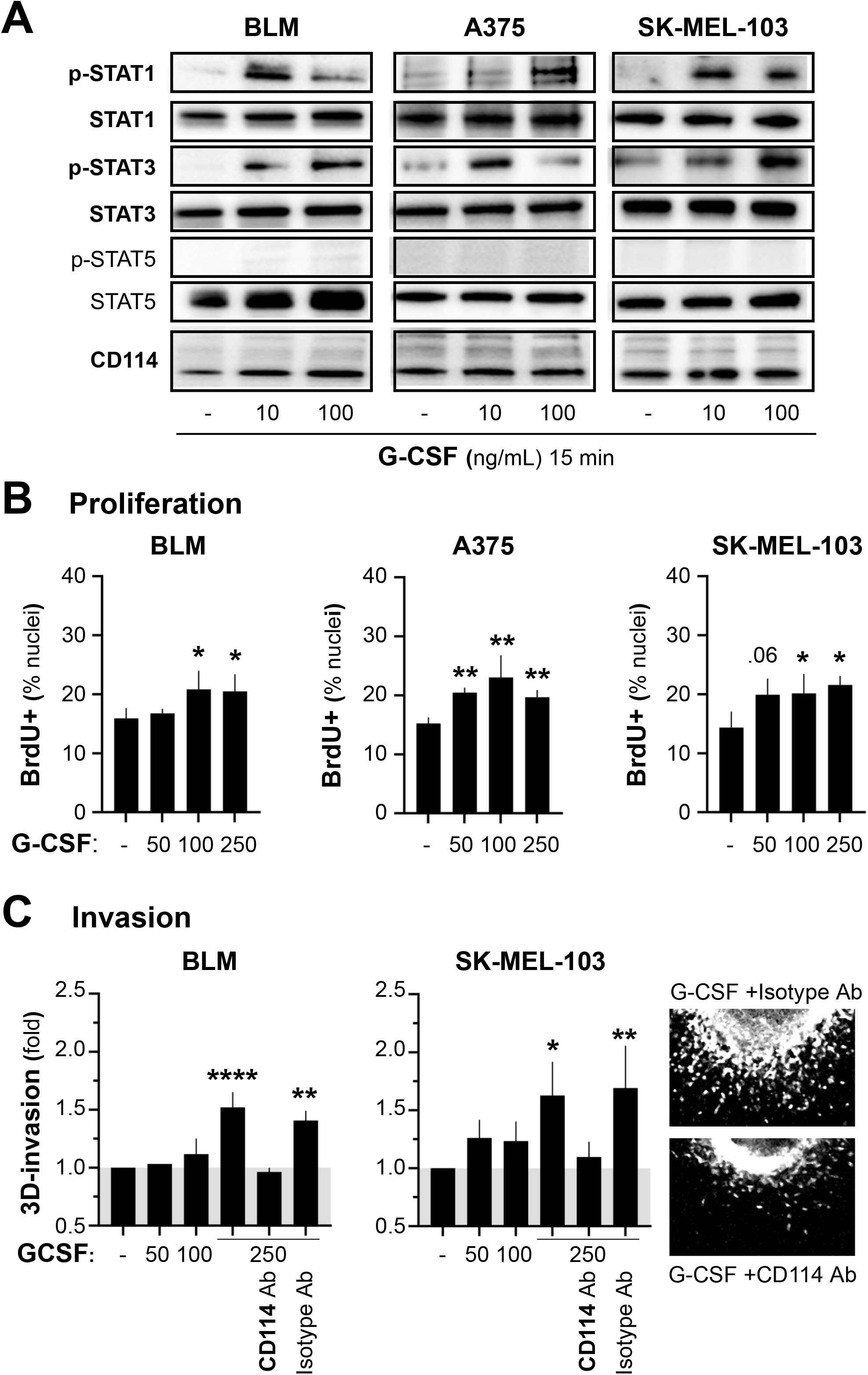
Direct effect of G-CSF on melanoma cell signaling and function. **(A)** Signaling kinetics in BLM, A375, and SK-MEL-103 melanoma cell lines whole lysates following stimulation with rhG-CSF at 10 and 100 ng/mL for 15 minutes. Immunoblots show phosphorylated STAT1 (Tyr701), STAT3 (Tyr705), STAT5A/B (Tyr694/699) and total forms of the proteins, alongside with CD114. Blots are representative of two independent experiments. **(B)** Proliferation assays for the three melanoma cell lines is serum-deprived media containing different concentrations of rhG-CSF (ng/mL). BrdU was pulsed for 1 hour and percentages of BrdU^+^ nuclei were quantified. Mean± SD values and significant differences to untreated controls, are shown (*p < 0.05; **p < 0.01; Mann– Whitney test). **(C)** BLM and SK-MEL-103 spheroids embedded in 3D collagen and allowed to invade for 72 hours in the presence of increasing concentrations of exogenous rhG-CSF (ng/mL), as indicated. Functionality of G-CSF receptors on melanoma cells was assessed using neutralizing antibodies against CD114 and its corresponding control isotype. Representative images of propidium iodide-stained BLM spheroids are shown. Mean ± SD values are represented and significant differences to untreated controls are indicated (*p < 0.05; **p < 0.01; ****p < 0.0001, one-way ANOVA and Tukey’s multiple comparison test). Functional assays (B, C) were repeated 2-5 times in duplicate.

### G-CSF and GM-CSF cooperate to produce a distinct macrophage differentiation state which enhances melanoma cell invasion

As already shown, monocytes infiltrating metastasizing melanoma tumors encounter a microenvironment specially enriched in G-CSF and GM-CSF, whereas those infiltrating non-metastasizing tumors are majorly exposed to constitutive M-CSF (Figure 4A). Thus, in order to determine the impact of G-CSF on monocytes, we treated them alone or in combination with GM-CSF and/or M-CSF in different assays. Expression of *CSF3R* was first confirmed by RNA-seq in monocytes and macrophages isolated from stage IV melanoma patients, as well as in monocytes from healthy donors and in vitro-primed macrophages. *CSF3R* transcripts were detected in all cell types, with particularly elevated expression in circulating monocytes from melanoma patients compared with healthy donors (Figure 4B). Interestingly, this upregulation was not observed for other hematopoietic myeloid receptors (Supplementary Figure S2A), and these patient-derived monocytes were enriched in a gene signature previously reported in G-CSF-mobilized monocytes [25] (Supplementary Figure S2B). These observations suggest a systemic deviation of hematopoiesis and priming of newly produced monocytes for subsequent G-CSF signaling at tumor level, which has been considered to play an important role in the tumor immunosuppressive milieu, metastasis and prognosis [26]. At the protein level, CD114 was expressed in monocytes and in vitro-derived macrophages (Figure 4C and Supplementary Figure S2C). Upon stimulation with rhG-CSF, phosphorylation of STAT1 and STAT3, but not STAT5, was induced in a dose-dependent manner within 15 minutes in monocytes. (Figure 4C). Given that the simultaneous expression of G-CSF and GM-CSF occurs within the metastasizing tumor microenvironment, we evaluated the effects of each cytokine individually as well as in combination. Notably, co-treatment with both cytokines induced concurrent activation of STAT1, STAT3, and STAT5 signaling pathways in monocytes (Figure 4D).

**Figure 4.**
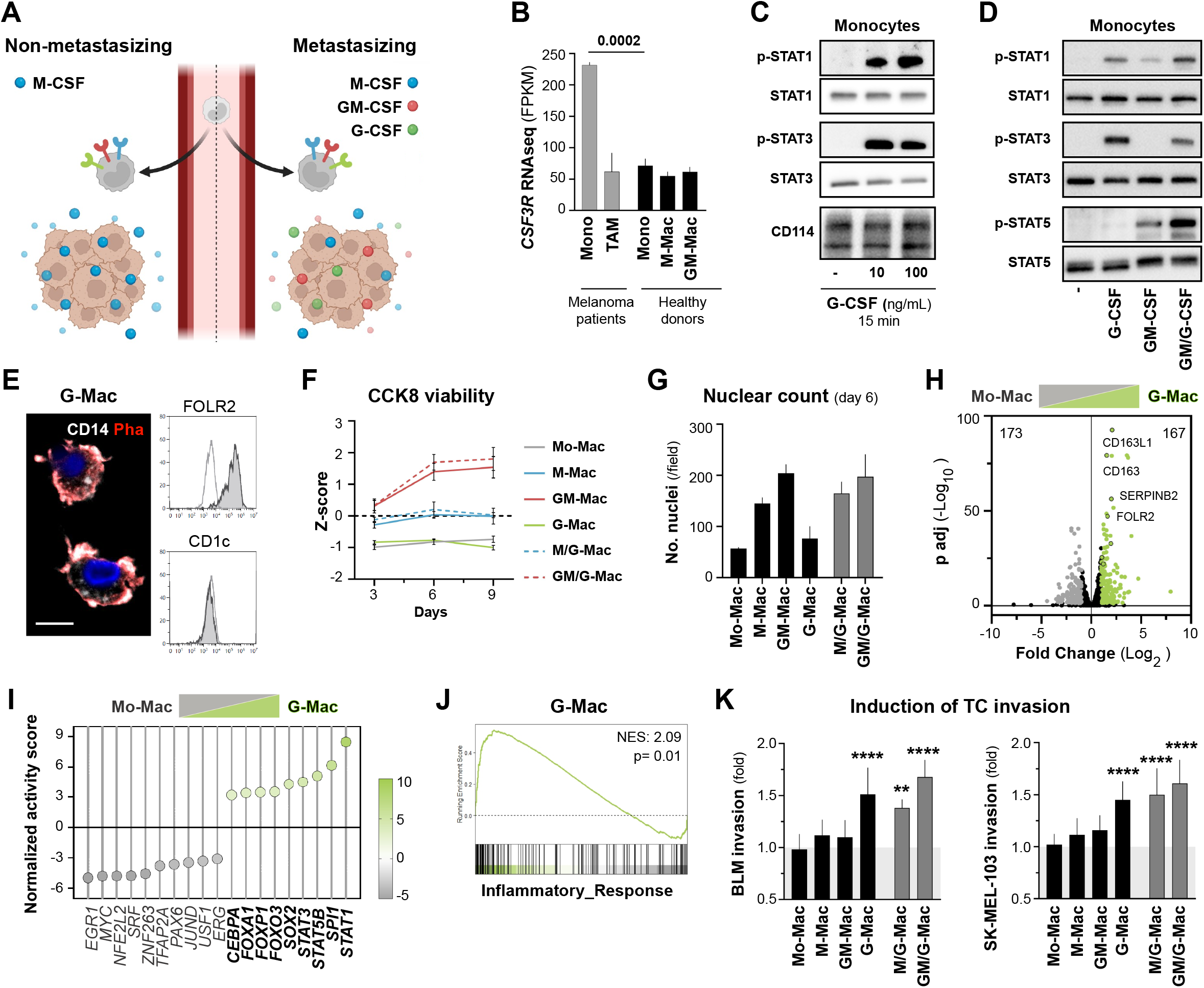
Direct effect of G-CSF on monocyte signaling and differentiation: pro-tumoral role of G-Macs. **(A)** Schematic representation of CSF expression in metastasizing and non-metastasizing primary melanomas. **(B)** *CSF3R* gene expression in monocytes and macrophages from healthy donors (n = 3), and monocytes or primary tumor TAMs from stage-IV melanoma patients (n = 3). Mean ± SEM values and unpaired t test analysis, are shown. **(C)** Signaling kinetics analyzed in fresh monocyte whole lysates following stimulation with rhG-CSF at 10 and 100 ng/mL for 15 minutes. Immunoblots show phosphorylated STAT1 (Tyr701), STAT3 (Tyr705), and total forms of the proteins, alongside with CD114. **(D)** G-CSF and GM-CSF signaling, alone or combined, analyzed in fresh monocyte whole lysates following stimulation with 10 ng/mL cytokine for 15 minutes. Immunoblots here include STAT5A/B (Tyr694/699). Blots are representative of 3 independent donors. **(E)** G-Macs (day 7) stained with anti-CD14 (white), Phalloidin (red), and Dapi (blue); or labelled for FOLR2 and CD1c as macrophage- and DC-lineage specific markers, respectively. Flow cytometry histograms are representative of 1 out of 5 donors (control, white; staining, grey). **(F)** Average cell viability and dehydrogenase activity of monocytes treated with indicated CSF combinations along 9 days. **(G)** Dapi-stained nuclear counts by field of indicated macrophages at the day 6 of treatment. F and G, Mean ± SD values are shown (n = 3 donors). **(H)** Volcano plot showing differential gene expression of monocyte-derived macrophages after 48 hours of G-CSF treatment (G-Macs) compared with untreated control (Mo-Macs, n = 3 donors). Significantly upregulated and downregulated genes are highlighted in green and gray, respectively (adjusted p value < 0.05; |log_2_FC| > 1). **(I)** Significant changes in transcription factor activities calculated with DoRothEA for ranked genes in G-Macs compared with Mo-Macs at 48 hours post-rhG-CSF treatment. Positive and negative activity scores are plotted in green and gray, respectively (adjusted p-value < 0.05). **(J)** Hallmark enrichment analysis (GSEA) of inflammatory response gene set for 48-hour G-Macs compared to Mo-Macs. Normalized enrichment score (NES) and adjusted p value are shown. **(K)** BLM and SK-MEL-103 spheroids embedded in 3D collagen and monitored for invasion over 72 hours in the presence of conditioned media generated from distinct CSF-differentiated macrophages, as indicated (n= 3-6 donors). Significant differences to the untreated control are indicated (**p < 0.01; ****p < 0.0001, one-way ANOVA and Dunnett’s multiple comparison test).

We next investigated whether prolonged exposure to G-CSF could drive monocyte differentiation toward a macrophage phenotype, analogous to the effects reported for M-CSF and GM-CSF. After seven days of stimulation, the resulting cells (G-Macs) expressed canonical macrophage markers (CD14^+^ FOLR2^+^ CD11c^+^ HLA-DR^+^ CD16^−^ CD15^−^ CD1c^−^) (Figure 4E and Supplementary Figure S2D). However, their viability was lower than that of alternatively (M-Macs) or classically (GM-Macs) primed macrophages and similar to non-stimulated monocytes (Mo-Macs), as shown by both metabolic activity and nuclear count assays (Figure 4F, G). Of note, the presence of G-CSF combined with either M-CSF or GM-CSF substantially enhanced cell viability during monocyte differentiation. Morphologically, G-Macs exhibited a rounder phenotype distinct from that of more flattened GM-Macs and M-Macs (Figure 4E; Supplementary Figure S2D), while their invasive capacity in 3D collagen matrices was intermediate between these two phenotypes (Supplementary Figure S2E). As early as 48 hours after G-CSF treatment, developing G-Macs were transcriptionally distinct from Mo-Macs, exhibiting a significant upregulation of canonical macrophage genes (i.e. *CD163, CD163L1, SERPINB2* and *FOLR2*; Figure 4H), while DoRothEA analysis revealed elevated transcriptional activity for STAT1, STAT3, and PU.1, among others (Figure 4I). Notably, G-Macs displayed a more inflammatory phenotype when compared with Mo-Macs, with significant enrichment of pathways involving IFN-γ, NF-κB and STAT3 signaling (Figure 4J; Supplementary Figure S2F).

Finally, conditioned media from all in vitro-–generated macrophage populations were assessed for their ability to induce tumor cell invasion. Only CM derived from G-Macs, or from macrophages differentiated with G-CSF in combination with M-CSF or GM-CSF, significantly enhanced the invasive capacity of BLM and SK-MEL-103 melanoma cells (p < 0.001; Figure 4K). These results collectively suggest that the converging pro-tumoral and pro-survival effects of G-CSF and GM-CSF cooperate to skew infiltrating monocytes toward a highly viable macrophage phenotype with enhanced capacity to promote melanoma cell invasion.

### G-Mac-conditioned melanoma cells exhibit higher lung colonization capability in vivo

As enhanced invasive capacity in vitro often correlates with increased metastatic potential in vivo, we established a lung colonization model in NSG mice in which melanoma cells must survive in the bloodstream, extravasate through lung vessels, and adapt to a new environment to form microscopically quantifiable metastases. Prior to tail-vein injection, melanoma cells were conditioned in vitro through co-culture with the generated macrophages to assess changes in their metastatic capacity in vivo (Figure 5A). A375 cells co-cultured with G-Macs—designated as A375(G-Mac)—exhibited significantly increased lung colonization compared to unconditioned or conditioned cells with Mo-Macs, as evidenced by a higher density and larger size of melanoma micrometastases (Figure 5B and 5C). This result was reproduced with the melanoma cell line BLM that also responded to G-Mac conditioning with increased in vivo spreading (Figure 5C). To further characterize these effects, we performed transcriptomic profiling of BLM cells following 24 hours of co-culture with either G-Macs or Mo-Macs. Comparative GSEA revealed enrichment of multiple pathways in G-Mac-conditioned melanoma cells, including those associated with invasiveness (i.e. TNF_NFKB, EPITHELIAL_MESENCHYMAL_TRANSITION, HYPOXIA and INFLAMMATORY_RESPONSE) [27] and proliferation (MYC_TARGETS_V2, G2M_CHECKPOINT, E2F_TARGETS) (Figure 5D). Remarkably, an increased lung colonization capacity was also observed upon conditioning of melanoma cells with GM/G-Macs (Figure 5C), which accurately reflects the G-CSF and GM-CSF co-expression profile observed in human primary tumors with metastatic potential.

**Figure 5.**
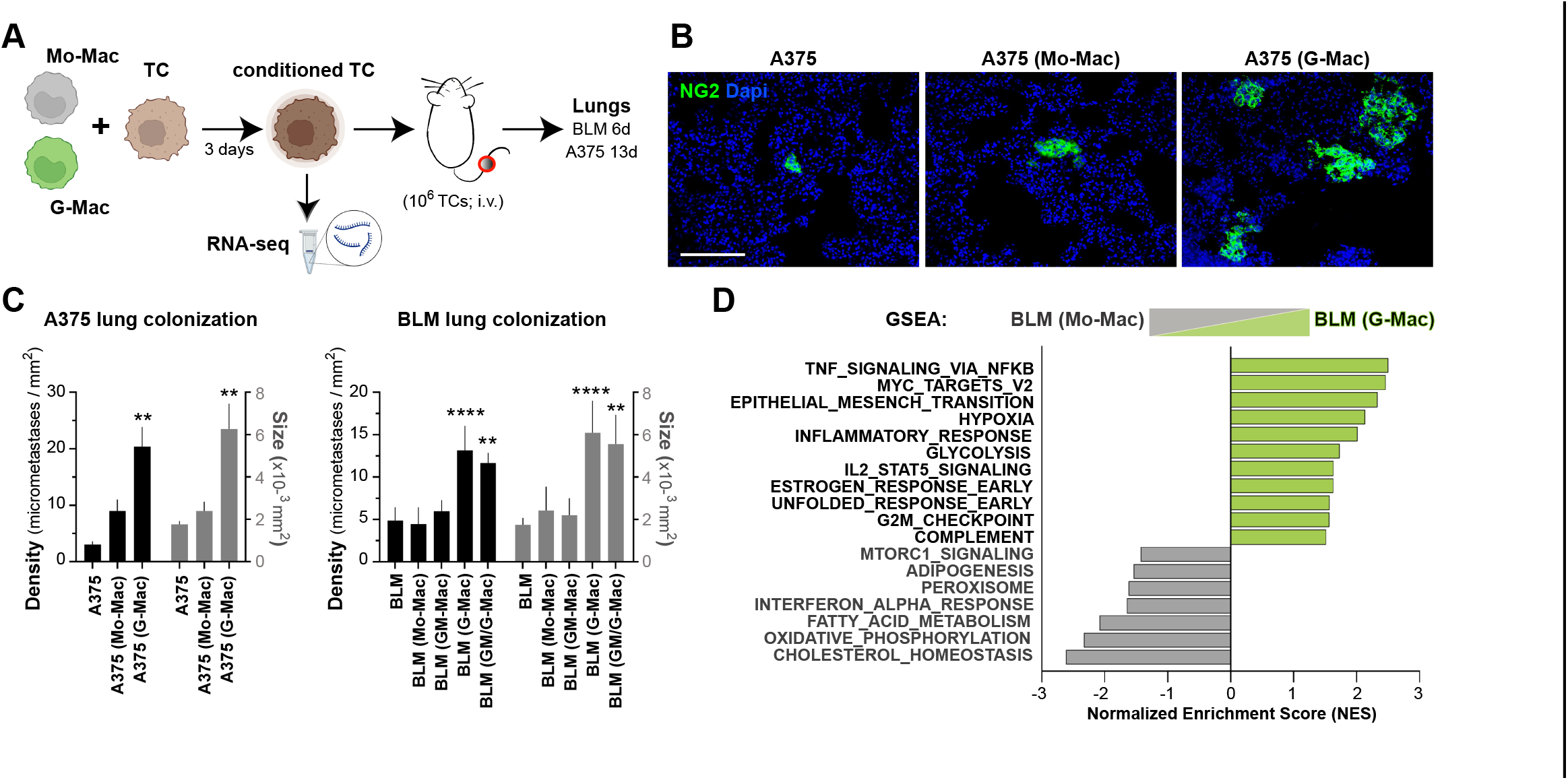
G-Mac-conditioned melanoma cells display enhanced lung colonization capacity in vivo. **(A)** Scheme of cell preparation and lung colonization model in immune-compromised NSG mice. **(B)**NG2^+^ stained melanoma colonies (green) are shown in representative lung sections from mice injected with A375 melanoma cells and A375 previously co-cultured with Mo-Macs [A375 (Mo-Mac)] or G-Macs [A375 (G-Mac)]. Dapi, blue. Scale bar, 300 µm. **(C)** Histograms showing quantifications of A375 and BLM lung colonies after 13 and 6 days of intravenous injection, respectively. Mean±SD density (number/mm^2^) and size (μm^2^) values of spontaneous melanoma micrometastases are shown (n= 3-6 mice per condition; 15-20 lung fields/mouse). Significant differences to the respective untreated control are indicated (**p < 0.01; ****p < 0.0001, one-way ANOVA and Dunnett’s multiple comparison test). **(D)** Hallmark enrichment analysis (GSEA) of BLM cells 24-hours conditioned by G-Macs [BLM (G-Mac), green] relative to Mo-Macs [BLM (Mo-Mac), grey]. NES of significantly upregulated gene sets are shown (adjusted p < 0.01).

## DISCUSION

The identification of reliable prognostic factors in human melanoma remains a major clinical challenge given its pronounced heterogeneity and the pivotal role of metastasis in patient outcome. Increasing evidence indicates that components of the TME critically influence melanoma behavior, highlighting immune cells and highly plastic TAMs as potential sources of prognostic information [1,4]. Using a melanoma–macrophage co-culture model, we observed multiple autocrine and paracrine signaling networks established between both cell types, particularly prominent when macrophages had been previously primed with GM-CSF, a cytokine predominantly expressed in tumors with metastatic potential [8]. Among these, we found the G-CSF/CD114 axis, which has been previously reported to modulate tumor initiation, progression, and metastasis either through direct pro-tumorigenic mechanisms or by modulating the immune infiltrate within the TME [13].

Analyses of publicly available transcriptomic datasets have revealed that the prognostic relevance of the *CSF3/CSF3R* axis is highly dependent on cancer type [28]. Elevated expression of both *CSF3* and *CSF3R* was significantly associated with increased TAM infiltration, advanced tumor grade, and poor patient prognosis in glioma [16,29,30], as well as invasion-related and immunomodulatory gene signatures in colorectal cancer [31,32]. In breast cancer, a G-CSF-responsive transcriptional signature was associated with reduced disease-free survival in luminal breast cancer, potentially through mechanisms involving MDSC accumulation [33]. Specifically in melanoma, *CSF3* expression has not been associated with patient prognosis [29], whereas elevated *CSF3R* expression was linked to improved clinical outcomes [28]. In contrast, our findings demonstrate by immunostaining that both G-CSF and its receptor CD114 are significantly enriched in TAMs and TCs from primary melanomas of patients who subsequently developed metastatic disease. These results provide a tissue-level protein perspective that contrasts with prior genomic data, resolving cell-type–specific expression patterns in stage II–IV tumors that may be obscured in bulk RNA sequencing approaches. The prognostic value of G-CSF was supported by its association with both reduced disease-free and overall survival. This finding differs from a report in breast cancer where G-CSF tissue immunodetection was associated with a favorable prognosis [34], but is consistent with other studies linking high G-CSF and CD114 protein expression to advanced tumor stage, metastasis, and poor patient survival across ovarian, cervical, colorectal, gastric, pancreatic, and esophageal cancers [35–41]. Moreover, we identified a positive correlation between G-CSF and GM-CSF expression, especially in poor outcome patients, similarly to a previous study in advanced-stage glioma [42]. Multivariate analysis revealed that G-CSF expression in TAMs exerts a dominant prognostic influence over GM-CSF, whereas the prognostic value of both cytokines in melanoma cells suggests an additive or potentially synergistic interaction role, where each ligand might contribute independently to disease progression.

Given that both TAMs and melanoma cells express G-CSF and its receptor, multiple potential signaling mechanisms may be engaged. Here, we demonstrate that both melanoma cell lines and human monocytes are directly responsive to G-CSF through activation of STAT1 and STAT3, but not STAT5. The G-CSF/STAT3 axis has been previously reported to mediate proliferation and migration of multiple tumor types [35,43–46]; accordingly, we found that rhG-CSF directly enhanced melanoma cell proliferation and collagen invasion in vitro. On the other hand, while the literature demonstrates that G-CSF exerts direct effects on monocytes in vitro primarily through selective STAT3 activation [47], leading to an attenuated cytokine response to LPS stimulation [48–51] or an impaired activation during dendritic cell (DC) differentiation [52,53], our data demonstrate that human monocytes simultaneously engage STAT1. This dual activation aligns with the transcriptional profile of G-CSF-differentiated macrophages (G-Macs), which exhibited significant enrichment in distinct inflammation and interferon-related gene signatures that may reflect a pro-inflammatory cell state.

Functionally, G-CSF alone did not promote monocyte survival but induced a secretory phenotype that enhanced melanoma cell invasion, consistent with previous results of Hollmén et al. where conditioned media from G-CSF-differentiated macrophages promoted breast cancer cell migration [54]. While M-CSF is constitutively expressed in primary melanoma tissues, GM-CSF and/or G-CSF are preferentially enriched in the TME of pro-metastatic lesions. To address this milieu complexity, we evaluated the combined effects of CSFs and found them to be additive, as STAT signaling pathways were simultaneously activated and the pro-survival function of GM-CSF complemented the pro-tumoral activity induced by G-CSF. Furthermore, melanoma cells conditioned by G-Macs exhibited transcriptomic and functional reprogramming, with an enrichment of pro-tumorigenic and invasive gene pathways in vitro and enhanced lung colonization in vivo. This observation underscores the indirect pro-tumoral effect of G-CSF through macrophage-tumor crosstalk, in accordance with a previous murine report where pro-tumorigenic macrophage activity was abrogated by G-CSF receptor knockout [55]. Collectively, the enrichment of the G-CSF/CD114 axis in melanoma cells and TAMs from tumors with metastatic potential supports the existence of an autocrine and paracrine signaling loop that directly promotes tumor growth and invasion while sustaining a reciprocal axis that favors melanoma aggressiveness.

In addition to our results in melanoma, growing evidence suggests that G-CSF within the tumor microenvironment of various cancer types promotes malignant progression and/or metastatic dissemination, ultimately contributing to poor prognosis and reduced patient survival [39,37]. Nevertheless, rhG-CSF is widely administered in cancer patients following myelosuppressive cytotoxic chemotherapy or during lymphodepleting regimens prior to adoptive T-cell therapies (TILs) in metastatic melanoma, even when its impact on therapeutic efficacy, as well as its role in preventing infectious complications, has not yet been systematically evaluated [56]. Notably, two retrospective studies have reported an association between rhG-CSF administration and distant organ metastases in patients with breast cancer and non–small cell lung cancer [57,58]. Moreover, G-CSF-producing tumors exhibited resistance to adoptive cell transfer immunotherapy in mice [31], and it has been recently reported that G-CSF administration attenuates chemo-immunotherapy efficacy in small-cell lung cancer patients [59]. These reports underscore the need for a careful evaluation of potential effects of rhG-CSF in tumor progression, and its use in melanoma patients should take into consideration its potential pro-metastatic role by shaping the tumor-macrophage crosstalk.

## CONCLUSION

Our study identifies the G-CSF/CD114 axis as a critical, underappreciated component of the melanoma TME with significant prognostic and functional implications. G-CSF and its receptor are enriched in both TAMs and melanoma cells from metastasizing tumors, and G-CSF expression correlates with reduced disease-free and overall survival. Mechanistically, this axis drives a bidirectional tumor–macrophage signaling network via STAT1/3 activation where it directly promotes melanoma proliferation and invasion, while indirectly shaping a pro-tumorigenic macrophage profile. Importantly, the additive effects of G-CSF and other CSFs suggest that combinatorial cytokine signaling more accurately reflects the biological complexity of metastasizing melanomas. Beyond its biological relevance, this work raises critical translational implications regarding the clinical use of rhG-CSF in oncology, and positions the G-CSF/CD114 pathway as a potential prognostic biomarker and a candidate therapeutic target aimed at disrupting tumor–macrophage crosstalk during melanoma progression.

## ACKNOWLEDGMENTS

We thank the staff of the statistical, flow cytometry, cell culture, medical image, UPMTA and animal facilities at the IiSGM for their expert technical assistance.

## ETHICS STATEMENT

The studies involving humans were approved by Comité de Ética de la Investigación con Medicamentos del Hospital General Universitario Gregorio Marañón (CEIM 20/18). The studies were conducted in accordance with the local legislation and institutional requirements, adhered to the Declaration of Helsinki. The participants provided their written informed consent to participate in this study.

## DATA AVAILABILITY STATEMENT

The dataset generated or used in this study are publicly available in Gene Expression Omnibus with accession numbers GSE171277, GSE242674 and GSE (pending).

## AUTHOR CONTRIBUTIONS

Conceptualization: ANV, PSM, RS; Data Curation: ANV, LBR, CBA, JAAI, VPB, RS; Formal Analysis: ANV, LBR, BLN, CBA, RS; Funding acquisition: PSM, RS; Investigation: ANV, LBR, BLN, CBA, AFL, RS; Methodology: ANV, LBR, BLN, CBA, AFL, RS; Project administration: ANV, RS; Resources: JAAI, VPB, PSM, RS; Supervision: RS; Validation: ANV, LBR, BLN, CBA, RS; Writing-Original Draft Preparation: ANV, RS; Writing-Review and Editing: PSM.

## LIST OF ABBREVIATIONS

BrdU: 5-bromo-2’-deoxyuridine
CD114 or *CSF3R*: granulocyte colony-stimulating factor receptor
CM: conditioned media
CSFs: colony-stimulating factors
DC: dendritic cell
DEGs: differentially expressed genes
DFS: disease-free survival
DoRothEA: Discriminant Regulon Expression Analysis
FCS: fetal calf serum
FFPE: formalin-fixed and paraffin-embedded
FPKM: fragments per kilobase of transcript per million mapped reads
G-CSF or *CSF3*: granulocyte colony-stimulating factor
GM-CSF: granulocyte-macrophage colony-stimulating factor
GSEA: gene set enrichment analysis
M-CSF: macrophage colony-stimulating factor
MDSCs: myeloid-derived suppressor cells
MFI: mean fluorescence intensity
MSigDB: Molecular Signatures Database
NSG mice: NOD/SCID/IL-2Rγ−/− mice
OS: overall survival
rh: recombinant human
RNA-seq: RNA sequencing
TAMs: tumor-associated macrophages
TCs: tumor cells
TME: tumor microenvironment

## LEGENDS

**Supplementary Figure S1.**
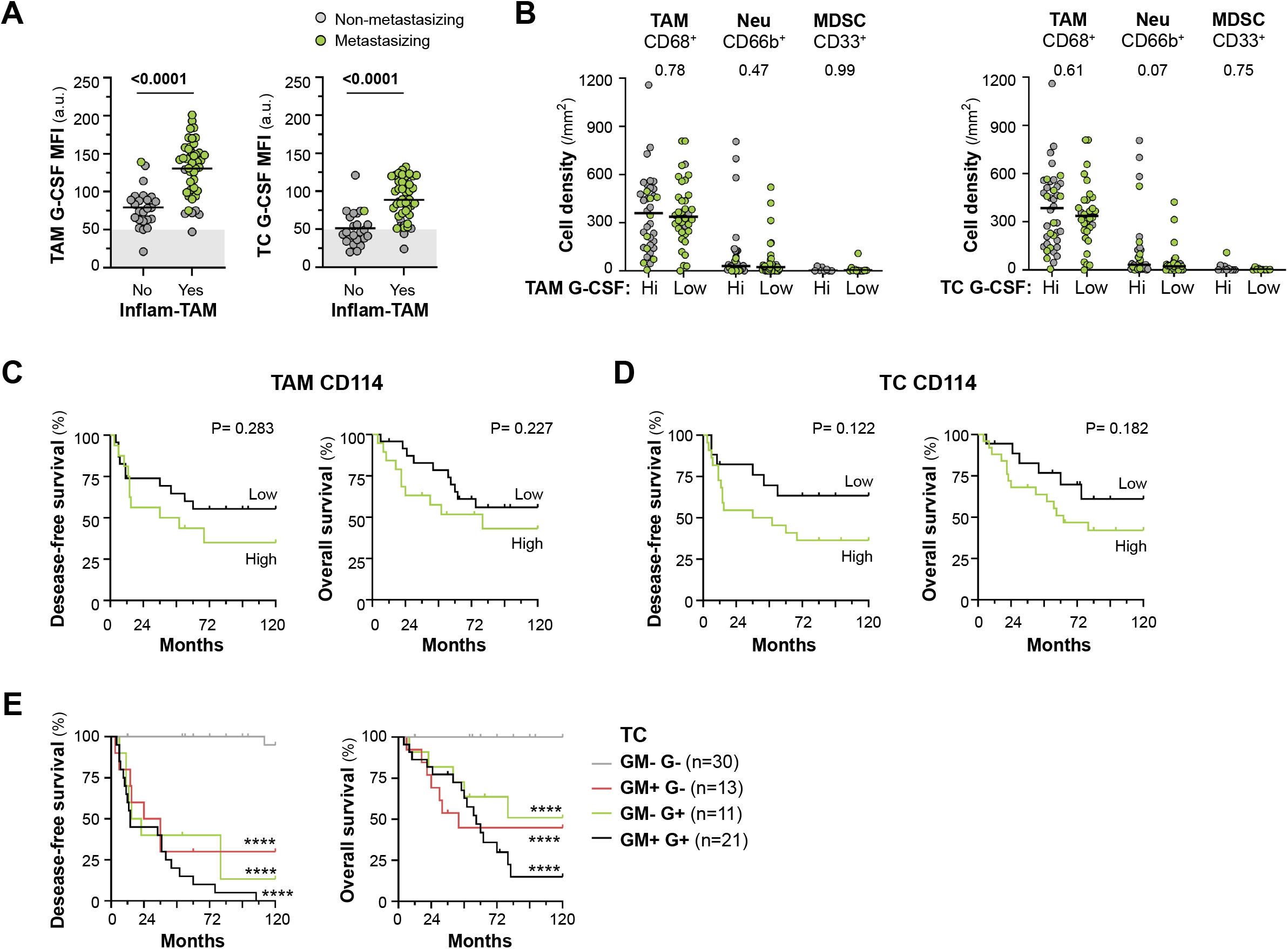
Primary melanoma tissue quantifications. **(A)** TAM and TC average G-CSF levels in primary melanomas populated or not by inflammatory cytokine-producing TAMs (Inflam-TAM), as previously classified [7]. MFI for each sample and Mann–Whitney p values, are shown. **(B)** Intratumoral cell density of CD68^+^ TAMs, CD66b^+^ neutrophils, and CD33^+^ MDSCs in primary melanomas classified by G-CSF expression in TAMs or TCs, as indicated. Mann-Whitney statistic p values are shown. In A and B, samples are colored as metastasizing or non-metastasizing, as noted. **(C, D)** Disease-free and overall survival 10-year Kaplan–Meier curves for CD114 expression in TAMs (C; cut-off value: 110 a.u.) and TCs (D; cut-off value: 150 a.u.). Log-rank p value*s* are shown. **(E)** Disease-free and overall survival 10-year Kaplan–Meier curves for GM-CSF and G-CSF expression in TCs, categorized by high (+) or low (-) expression of both cytokines, as indicated. Log-rank p value*s* relative to double negative samples are shown (****p < 0.0001).

**Supplementary Figure S2.**
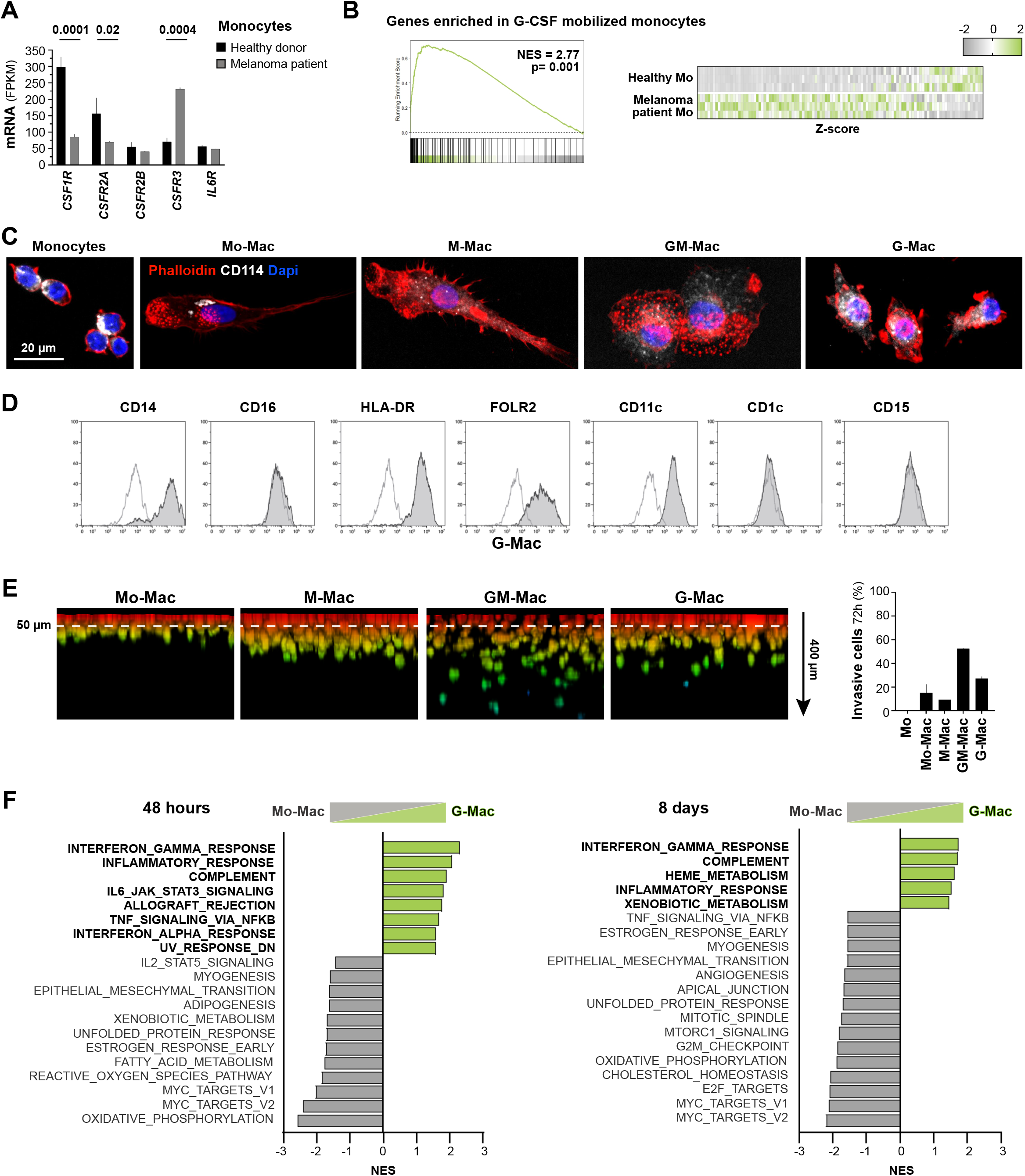
Macrophage profiles generated in vitro from blood monocytes. **(A)** Gene expression of indicated hematopoietic receptors as determined by RNA-seq on CD14^+^ monocytes isolated from healthy donors and stage-IV melanoma patients. Mann-Whitney statistic p values are shown (n = 3). **(B)** Expression heatmap and enrichment analysis of genes upregulated in G-CSF-mobilized monocytes [25] for our melanoma patient-derived CD14^+^ monocytes in comparison with healthy donors (n = 3). **(C)** Staining of monocytes (2 hours after isolation) and CSF-differentiated macrophages (Mo-Macs, M-Macs, GM-Macs, and G-Macs) after 7 days. Cells were grown on PLL-covered slides and stained for F-Actin (Phalloidin, red), CD114 (white), and Dapi (blue) after 0.5% Triton-X100 permeabilization. Scale bar = 20 µm. **(D)** Surface staining of G-Macs with indicated myeloid markers. Flow cytometry histograms are representative of 1 out of 5 donors (control, white; staining, grey). **(E)** Macrophage 3D-collagen invasion for 72 hours in 1% FCS medium. Representative lateral projections and percentages of invasive cells (> 50 µm) are shown (n = 3 donors). **(F)** Hallmark enrichment analysis (GSEA) of G-Macs (green) after 48 hours (left) or 8 days (right) of rhG-CSF-differentiation in comparison to untreated Mo-Macs (gray). NES of significantly upregulated gene sets are shown (adjusted p < 0.01).

